# Neighboring colonies influence uptake of thermotolerant endosymbionts in threatened Caribbean coral recruits

**DOI:** 10.1101/2020.11.02.360941

**Authors:** Olivia M. Williamson, Corinne E. Allen, Dana E. Williams, Matthew W. Johnson, Margaret W. Miller, Andrew C. Baker

## Abstract

Intervention strategies to enhance coral resilience include manipulating the association between corals and their algal endosymbionts. While hosting thermotolerant *Durusdinium trenchii* can increase bleaching thresholds in adults, its effects remain largely unknown during the early life stages of Caribbean corals. Here, we tested if *Orbicella faveolata* recruits could establish symbiosis with *D. trenchii* supplied by nearby “donor” colonies and examined the resulting ecological trade-offs to evaluate early Symbiodiniaceae manipulation as a scalable tool for reef restoration. We exposed aposymbiotic recruits to 29°C or 31°C and to fragments of either *Montastraea cavernosa* (containing *Cladocopium*) or *Siderastrea siderea* (containing *D. trenchii*). After 60 days, recruits reared with *D. trenchii* donors hosted nearly three times more *D. trenchii* than those with *Cladocopium* donors, suggesting that recruits can acquire Symbiodiniaceae from nearby corals of different species. Temperature did not affect *D. trenchii* uptake. Next, donor colonies were removed and surviving recruits were maintained for three months at ambient temperatures, after which a subset was exposed to a 60-day heat stress trial. Recruits previously reared at 31°C survived twice as long at 34°C as those reared at 29°C, suggesting that pre-exposure to heat can prime recruits to withstand future thermal stress. In addition, recruits hosting primarily *D. trenchii* survived twice as long at 34°C as those hosting little or no *D. trenchii*. However, the proportion of *D. trenchii* hosted was negatively correlated with polyp size and symbiont density, indicating a trade-off between growth (of both host and symbiont) and heat tolerance. These findings suggest that, while donor colonies may be effective sources for seeding coral recruits with thermotolerant symbionts, practitioners will need to balance the likely benefits and costs of these approaches when designing restoration strategies.

## Introduction

Scleractinian corals are ecosystem engineers, building reefs that support one quarter of all marine biodiversity and contributing billions of dollars to global economies annually through fisheries, tourism, and coastline protection (Jones et al. 1994; Wells et al. 2006). However, coral reefs are declining rapidly as multiple stressors outpace their natural capacity to evolve and threaten the critical ecosystem services they provide (Hughes & Connell 1999; Hughes et al. 2003; Pandolfi et al. 2003; Wild et al. 2011). A major cause of recent decline is coral bleaching (Hughes et al. 2017), a phenomenon wherein stressful environmental conditions destabilize the partnership between corals and Symbiodiniaceae, a diverse family of largely symbiotic dinoflagellate algae (LaJeunesse et al. 2018), prompting ejection of the algae by the coral hosts (Knowlton & Rohwer 2003; Weis 2008). However, some corals survive these bleaching episodes, recovering or even maintaining their symbioses while neighboring colonies perish (Baker et al. 2004; LaJeunesse et al. 2010; Cunning et al. 2016). One factor contributing to this resilience is the type(s) of Symbiodiniaceae hosted within the corals; when stressful conditions subside, bleached corals can reacquire algae (Jones & Yellowlees 1997; Boulotte et al. 2016; Cunning et al. 2016). Termed ‘symbiont shuffling’ (Baker 2003), this process involves shifts in the relative abundance of different symbiont types within the host, which can result in replacement of the numerically dominant symbiont and alter the community function (Buddemeier & Fautin 1993; Toller et al. 2001; Baker 2001; Stat et al. 2008).

Many members of the Symbiodiniaceae genus *Durusdinium* are particularly tolerant to thermal stress (Glynn et al. 2001; Baker et al. 2004; Rowan 2004) and may prove critical for the persistence of reefs. Most prevalent after bleaching events or on reefs characterized by extreme or highly variable conditions (Baker et al. 2004; Fabricius et al. 2004; LaJeunesse et al. 2009, 2010), *Durusdinium* is widely distributed (albeit at generally low abundance) among locations and host species worldwide (Silverstein et al. 2012). During recent marine heatwaves, corals hosting this genus have maintained their symbioses while nearby colonies bleached (Glynn et al. 2001; Baker et al. 2004; LaJeunesse et al. 2009; Kemp et al. 2014), and many bleached colonies have recovered with predominantly *Durusdinium*, increasing the holobiont’s subsequent bleaching threshold by ∼1-2°C (Berkelmans and van Oppen 2006; Silverstein et al. 2015). Thus, symbiont shuffling, and particularly relative increases in *Durusdinium trenchii*, represents a mechanism of rapid ecological acclimatization and a potential tool for researchers to enhance coral heat tolerance (Baker et al. 2004; Silverstein et al. 2015; Cunning et al. 2018; National Academies 2019).

However, despite its thermotolerance benefits, hosting *D. trenchii* may be accompanied by physiological trade-offs that impact whether symbiont shuffling is sustainable at the ecosystem scale. Studies have found reduced calcification, growth rates, lipid stores, and egg size in corals hosting *D. trenchii* under non-stressful temperatures (Little et al. 2004; Jones & Berkelmans 2011; Poquita-Du et al. 2020), suggesting that, while this symbiont increases heat tolerance, it may compromise long-term reef recovery (Ortiz et al. 2013; Pettay et al. 2015). Consequently, we must consider both the benefits and trade-offs of *D. trenchii* in the context of predicted stress exposure as we assess approaches to increase reef resilience.

While laboratory studies have manipulated algal partners in adult Caribbean corals to enhance stress tolerance (Silverstein et al. 2015; Cunning et al. 2016, 2018), the capacity of Caribbean coral juveniles to acquire *D. trenchii* and the eco-physiological trade-offs of this association are not well understood. Many corals are broadcast spawners, beginning life as aposymbiotic larvae that acquire Symbiodiniaceae “horizontally” from their environment (Harrison & Wallace 1990; Coffroth et al. 2006). A recruit’s algal partners influence fitness (Mieog et al. 2009), and thus early establishment of environmentally-appropriate symbioses may enhance survival (Chamberland et al. 2017; Quigley et al. 2017a). Various processes may govern symbiont acquisition and selection during early ontogeny (Little et al. 2004), but few studies have tracked the biotic and abiotic factors shaping symbiotic partnerships in young Caribbean corals. Indeed, Quigley et al. (2018) urged that research be directed at optimizing natural and artificial Symbiodiniaceae delivery to boost juvenile survival.

This study investigated whether temperature and/or neighboring adult corals could enhance initial *D. trenchii* uptake in juvenile *Orbicella faveolata*, an important Caribbean reef-builder listed as threatened under the Endangered Species Act. First, we tested the hypothesis that elevated temperature and/or proximity of *D. trenchii* “donor” colonies increases *D. trenchii* abundance in recruits during symbiosis establishment. Next, we examined the physiological trade-offs, such as polyp size and algal cell density, of hosting *D. trenchii*. Finally, we tested if symbiont community and/or previous heat exposure impacted recruit survival during a heat stress trial.

## Methods

### Larval rearing and settlement

This experiment utilized newly-settled *Orbicella faveolata* recruits collected as gametes during a broadcast spawning event six days after the full moon in August 2018 from Horseshoe Reef in Key Largo, FL (25.1388°N, 80.2950°W). This location was chosen because multiple *O. faveolata* colonies were previously mapped, genotyped, and observed to spawn (Miller et al., 2018; Fisch et al., 2019). Gamete bundles were collected from eight colonies in conical mesh nets, gathering in removable 50 mL Falcon tubes at the nets’ apices. Once ∼20% filled with bundles, tubes were removed, capped, and brought to the boat. Gametes from all parents were pooled in 2-liter plastic containers, then immediately diluted with filtered seawater (FSW) as bundles began breaking apart to reach a sperm concentration of ∼10^6^ cells/mL following Hagedorn et al. (2009). Batches of mixed gametes were transported to a field lab in Key Largo, FL, where larvae were maintained in FSW for four days as they developed. Fresh FSW was provided and dead larvae were removed several times daily.

Four days post-fertilization, ∼4,000 larvae were transported to laboratories at the Rosenstiel School of Marine and Atmospheric Science, where they were placed in one-micron FSW and supplied with 108 ceramic plugs (2.5 cm diameter) to facilitate settlement. Plugs had one-mm grooves in the top and were previously preconditioned at Emerald Reef (near Miami, FL) for three weeks to develop a “reef-like” biofilm (Hadfield 2011). After two days, settlers were counted under a microscope. To maintain consistent light conditions, only recruits on the top face of plugs were tracked and sampled for the subsequent experiment (n=1,595).

### Experiment 1: Symbiodiniaceae uptake

Plugs were distributed randomly into 12 new 2.5-gallon aquaria, so that nine plugs started in each with an approximately even number of settlers (∼133 per aquarium). Aquaria were supplied with one-micron FSW (fine enough to exclude Symbiodiniaceae cells) and immersed in one of two 50-gallon seawater tanks to maintain temperatures, one held at 31°C and the other at 29°C. The maximum monthly mean (MMM) temperature on Horseshoe Reef in Key Largo, FL is ∼30°C (according to Pathfinder 5.0 and Coral Reef Watch – R. van Hooidonk, pers. comm.), so these treatments were within the scope of what newly-settled recruits might naturally experience. The 31°C treatment was designed to exert only mild heat stress on recruits without high mortality. Light (150–300 μmol quanta m−2 s−1, measured by an Apogee Underwater Quantum PAR Meter MQ□210) was maintained on a 12 hour light–dark cycle using 400 W metal halide pendant lights (IceCap Inc., USA). Irradiance and temperature were recorded every 20 minutes with a HOBO Pendant® data logger (Onset Computer Corporation MX2202). One ∼6 cm^2^ fragment of *Siderastrea siderea* was placed into half of the aquaria at each temperature. These fragments originated from a single colony and were known to host >95% *D. trenchii*. One fragment of *Montastraea cavernosa* hosting >95% *Cladocopium* was placed in each of the other six aquaria. One 4W submersible pump (VicTsing CAAA3-HG16) was placed into each aquarium to distribute heat and symbiont cells. With two temperatures and two symbiont sources, the experiment consisted of four treatments, each conducted in triplicate (Fig. 1).

**Figure 1:**
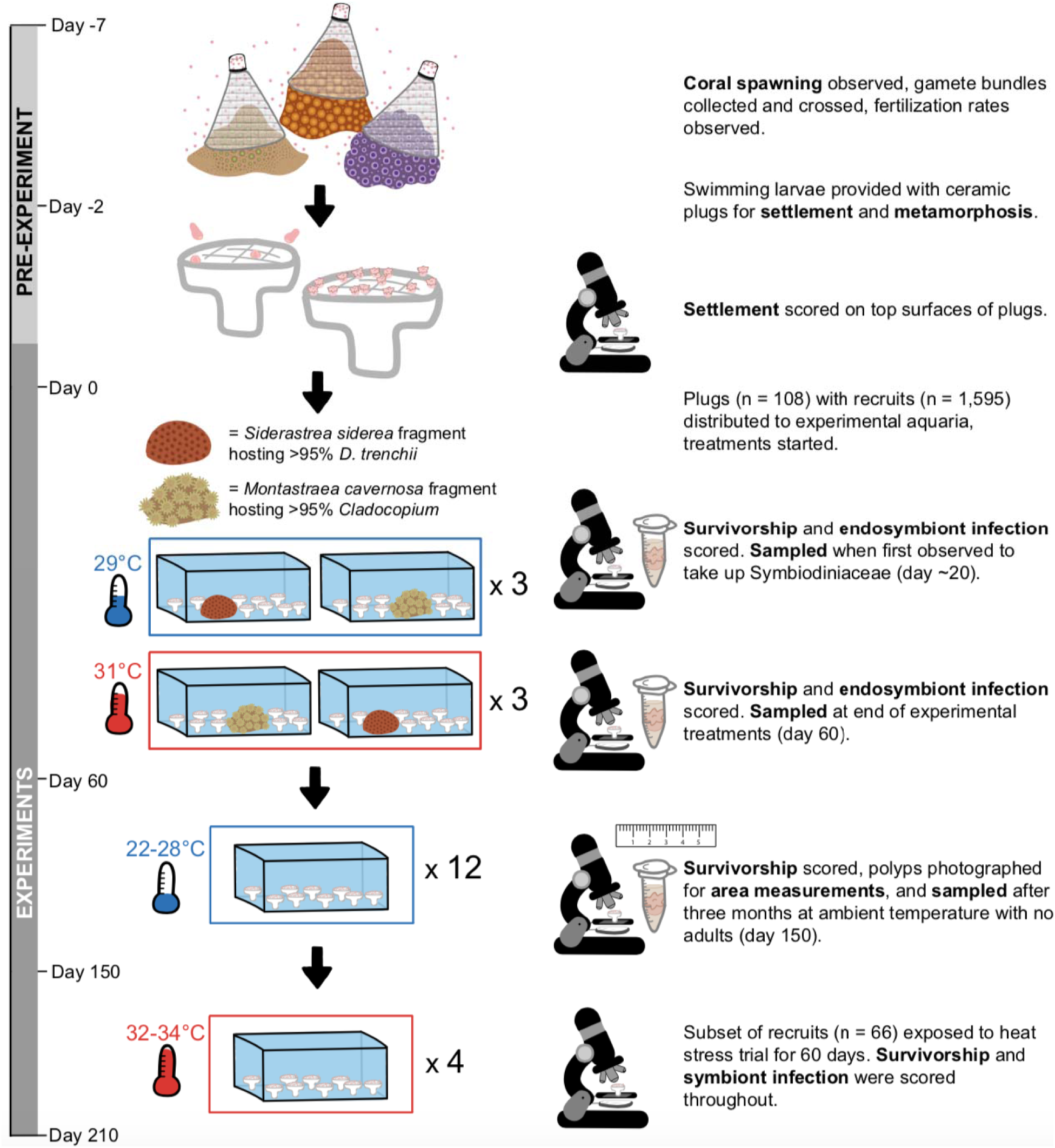
Sequence of experimental treatments and sampling events described in this study.

Recruits were maintained in their respective treatments for 60 days. FSW was replaced every other day, and algae were manually removed from plugs regularly to prevent overgrowth and/or competition with recruits. To minimize differences in light and flow, aquaria were shuffled within their temperature baths and plugs were shuffled within aquaria weekly.

### Observation and tissue sampling

At least once per week, a dissecting microscope was used to count the number of: (1) surviving recruits in all aquaria and (2) recruits visibly infected with symbionts (Fig. 2a, 2b). When infection was first observed in all aquaria (day 21) and again at the end of the 60 days in their treatments, three to five recruits from each aquarium were sacrificed using a razor blade (Fig. 1; Fig. 2d). To ensure consistency of sampling, only individual polyps not clumped with others were sacrificed (Fig. 2c). Sacrificed recruits were placed in individual 1.5-mL microcentrifuge tubes with 200 μL of 1% SDS + DNAB and incubated at 65°C for one hour. Genomic DNA was extracted from SDS archives following modified organic extraction methods (Rowan and Powers, 1991; Baker & Cunning, 2016).

**Figure 2:**
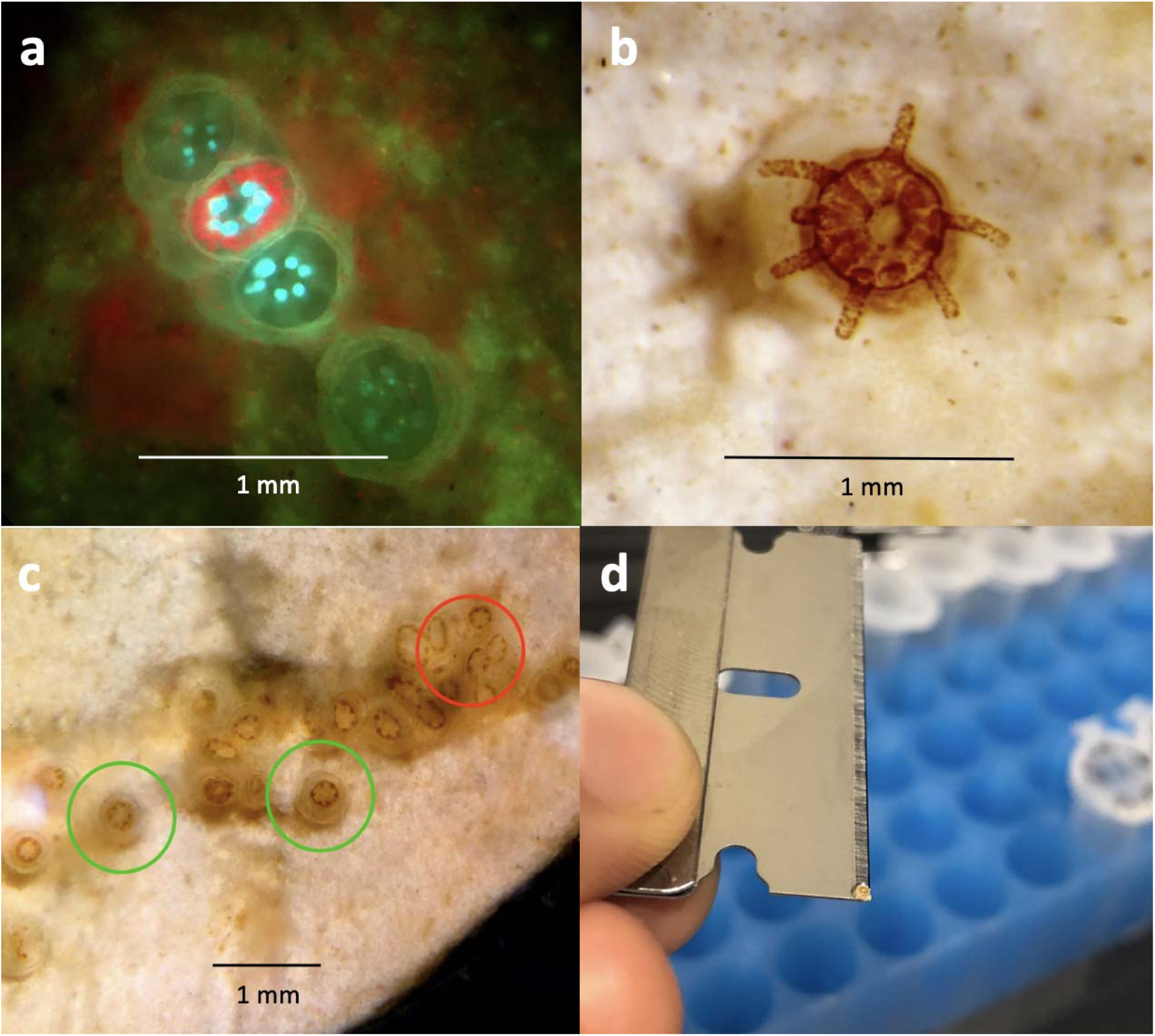
(a) Symbiodiniaceae infection was scored visually, aided by violet light filters to view red chlorophyll fluorescence in symbiont cells. (b) A one-month-old *O. faveolata* recruit infected with Symbiodiniaceae. **c)** Recruits circled in green are examples of individual, well-separated polyps, which were chosen for sampling. Recruits circled in red are clumped together without clear separation, and were not chosen for sampling. **d)** Individual recruits were sacrificed using a razor blade.

### Growth and symbiont density

After 60 days, donor fragments were removed from aquaria. Temperatures in aquaria at 31°C were reduced by 1°C per day to reach 27°C (the ambient temperature of incoming Bear Cut, FL, seawater at the time), while those at 29°C were reduced by ∼0.5°C per day. After this point temperature tracked local seasonal fluctuations, eventually reaching 22°C three months later (January 2019).

Five months (150 days) after settlement, a random sample of recruits from each of the four original treatments were photographed under a dissecting microscope with QCapture Suite Plus. Only individual, well-separated polyps were photographed to maximize accuracy of area measurements (Fig. 2c). ImageJ was used to calculate recruit skeletal area in mm^2^. R was used to create generalized linear models (GLMs) to compare recruit area by experimental temperature and symbiont sources. A subset of the photographed recruits were then sacrificed for DNA extraction and qPCR, to calculate symbiont identity and density at the time of growth measurements.

### Experiment 2: Heat stress trial

To assess whether: (1) previous exposure to elevated temperature and/or (2) *D. trenchii* dominance increased tolerance to future thermal stress, a subset of recruits were exposed to a heat stress trial in late January 2019 (day 150), three months after the end of treatments from Experiment 1. Three recruits per aquarium (nine per treatment, 36 total) were sampled to characterize Symbiodiniaceae communities in recruits from each of the 12 aquaria at the start of heat stress. However, since the small size of recruits prevented us from sampling without sacrificing them, we could not directly sample Symbiodiniaceae in individual recruits used in the heat stress trial. For analysis, aquaria were therefore split into three categories based on the mean proportion of *Durusdinium* hosted on day 150 (“low” = <0.25, “intermediate” = 0.25-0.75, “high” = >0.75; Fig. 7b).

Then, two or three plugs from each aquarium were placed into new aquaria with one-micron FSW, temperature was increased from 22°C to 28°C over six days, and then from 28°C to 32°C over 48 hours. At the start of this trial, all recruits (n=66) were infected with symbionts. About half the recruits (n=32) had been reared at 31°C during Experiment 1 and were thus pre-exposed to mild heat stress, while the other half (n=34) were reared at 29°C and naïve to heat stress. Temperature was maintained at 32°C for ten days, then raised to 33°C for ten days, then raised to 34°C for 40 days. Every two to five days, recruits were examined using a fluorescence microscope and scored as “healthy”, “pale”, “bleached”, or “dead” (Fig. 7, S2, S3). GLMs were created in R to test how prior heat exposure and proportion *Durusdinium* impacted survival and bleaching during heat stress.

### Symbiodiniaceae identification and quantification

Quantitative PCR (qPCR) assays were used to identify Symbiodiniaceae to genus level and quantify symbiont-to-host (S:H) cell ratios for each recruit sampled. Since *O. faveolata* commonly hosts members of the genera *Symbiodinium, Breviolum, Cladocopium*, and *Durusdinium* (Kemp et al., 2015), assays targeting specific actin loci for each genus were performed using a QuantStudio 3 Real-Time PCR Instrument (Applied Biosystems, USA). Assays for *O. faveolata, Symbiodinium*, and *Breviolum* followed reactions described in Cunning and Baker (2013), whereas *Cladocopium* and *Durusdinium* assays were multiplexed and conducted as described in Cunning et al. (2015a). The StepOneR software package in R was used to quality-filter assay results and calculate relative abundance of each symbiont genus and S:H cell ratios. The Kaplan-Meier estimate was used to calculate the fraction of recruits in each treatment living after a given amount of time (Goel et al., 2010). GLMs were created to compare the effects of each experimental factor and their interactions on survivorship, symbiont infection rates, symbiont community composition, and S:H cell ratios over time.

## Results

### Experiment 1: Symbiodiniaceae uptake

#### Survivorship and Symbiodiniaceae infection

Recruit survivorship did not vary significantly among treatments. In all treatments, the proportion of recruits visibly infected with Symbiodiniaceae increased over time (Fig. 3). Differences in proportion of infected recruits were driven primarily by temperature (p < 0.0001) and to a lesser extent by donor symbiont type (p = 0.048). A significantly higher proportion of recruits reared at 29°C were infected with symbionts compared with those reared at 31°C, which were delayed in their infection by ∼5-20 days at all time points after 14 days (Fig. 3). At 43 days at 31°C, a significantly higher proportion of recruits reared with *D. trenchii* donors were visibly infected compared to those with *Cladocopium* donors (p = 0.005).

**Figure 3:**
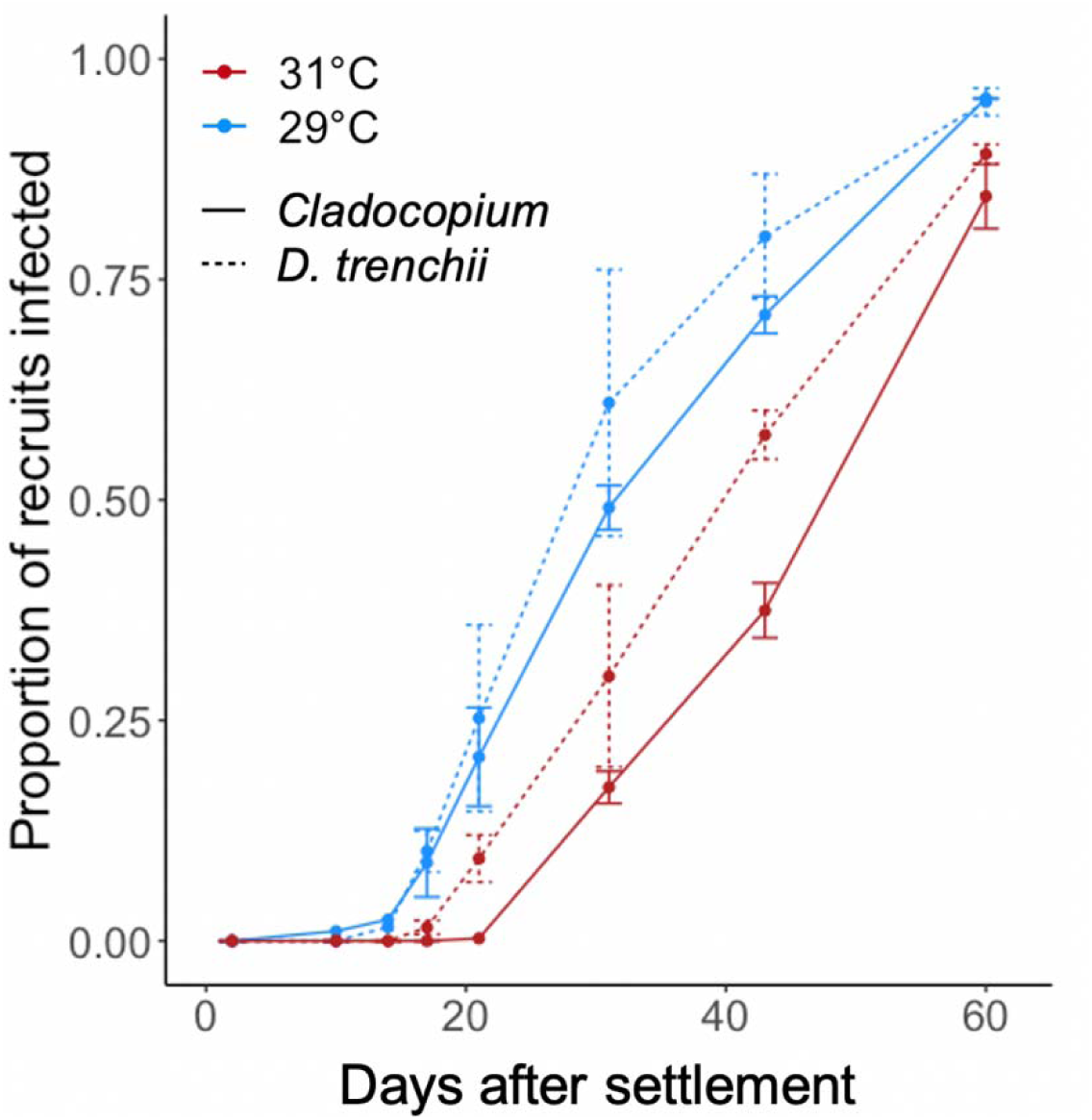
Symbiodiniaceae infection in recruits varied with temperature (p < 0.0001) and symbiont treatments (p = 0.048). Error bars represent +/- SEM.

#### Symbiodiniaceae identity and abundance

Recruits in all treatments hosted *Symbiodinium, Breviolum*, and *Durusdinium*, but *Cladocopium* was not detected. Overall, *D. trenchii* was found to be dominant (>90% of symbiont community) or co-dominant (10 – 90% of community) in 58.3% of recruits after 60 days. The proportion of *D. trenchii* in recruits was significantly related to donor symbiont type but not to temperature (Fig. 4). Recruits raised with *D. trenchii*-dominated colonies hosted higher proportions of *D. trenchii* (48.3±34.9%) than in those reared near *Cladocopium*-dominated colonies (17.8±24.9%) (p = 0.03).

**Figure 4:**
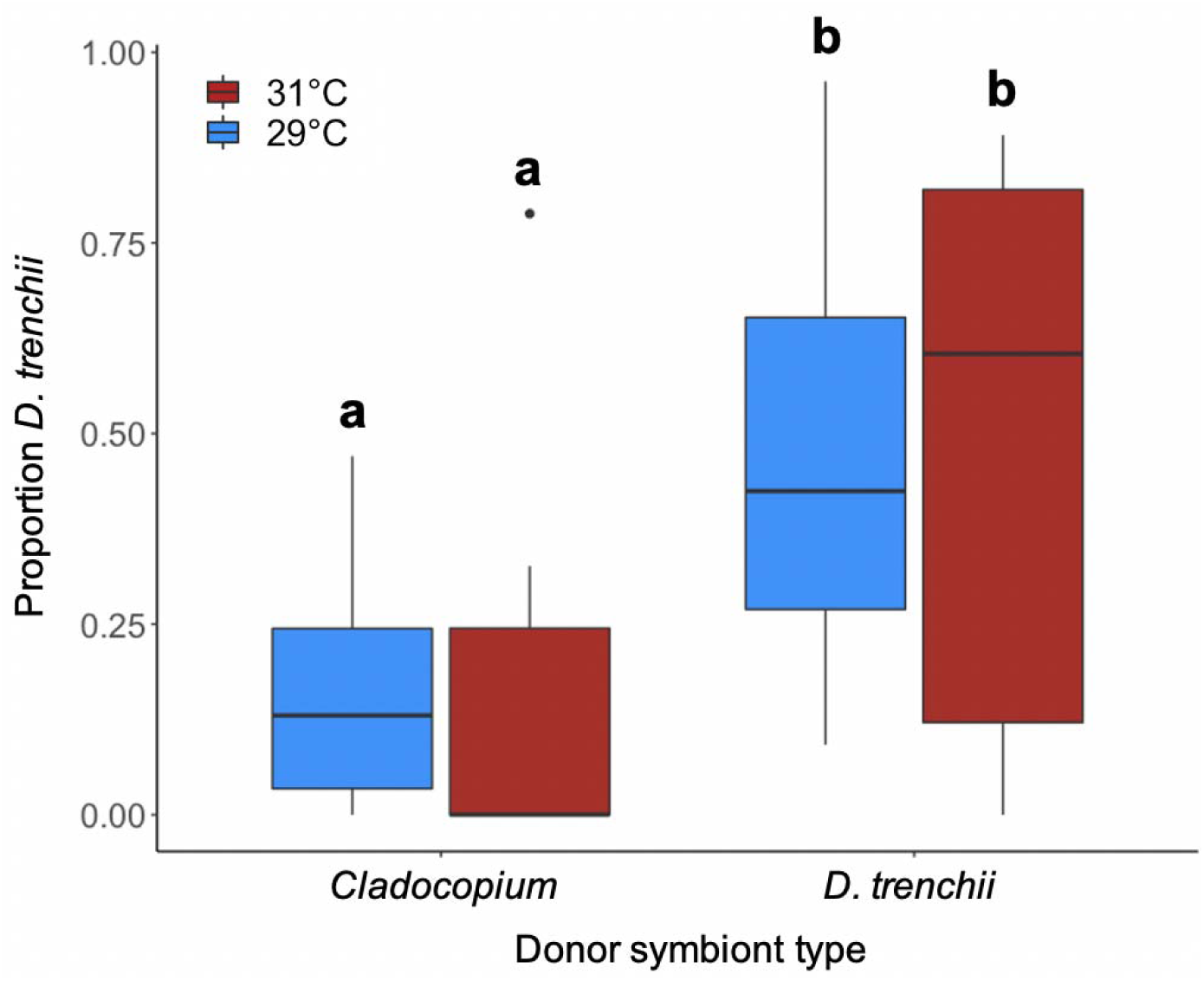
Proportion of *D. trenchii* in recruits reared near *D. trenchii* donors and *Cladocopium* donors after 60 days.

On average, S:H cell ratios increased in recruits with age but varied with symbiont community composition (Fig. 5) and showed significant interactions with time (p < 0.001). For recruits <150 days old, log S:H cell ratio decreased with increasing proportion of *Symbiodinium*, while in recruits >150 days old the opposite trend was observed (Fig. 5a). In recruits >14 days old, as proportion of *Breviolum* increased, the log S:H cell ratio also increased (p < 0.001, Fig. 5b). Conversely, as proportion of *D. trenchii* increased in recruits >14 days old, the log S:H cell ratio decreased (p < 0.001, Fig. 5c).

**Figure 5:**
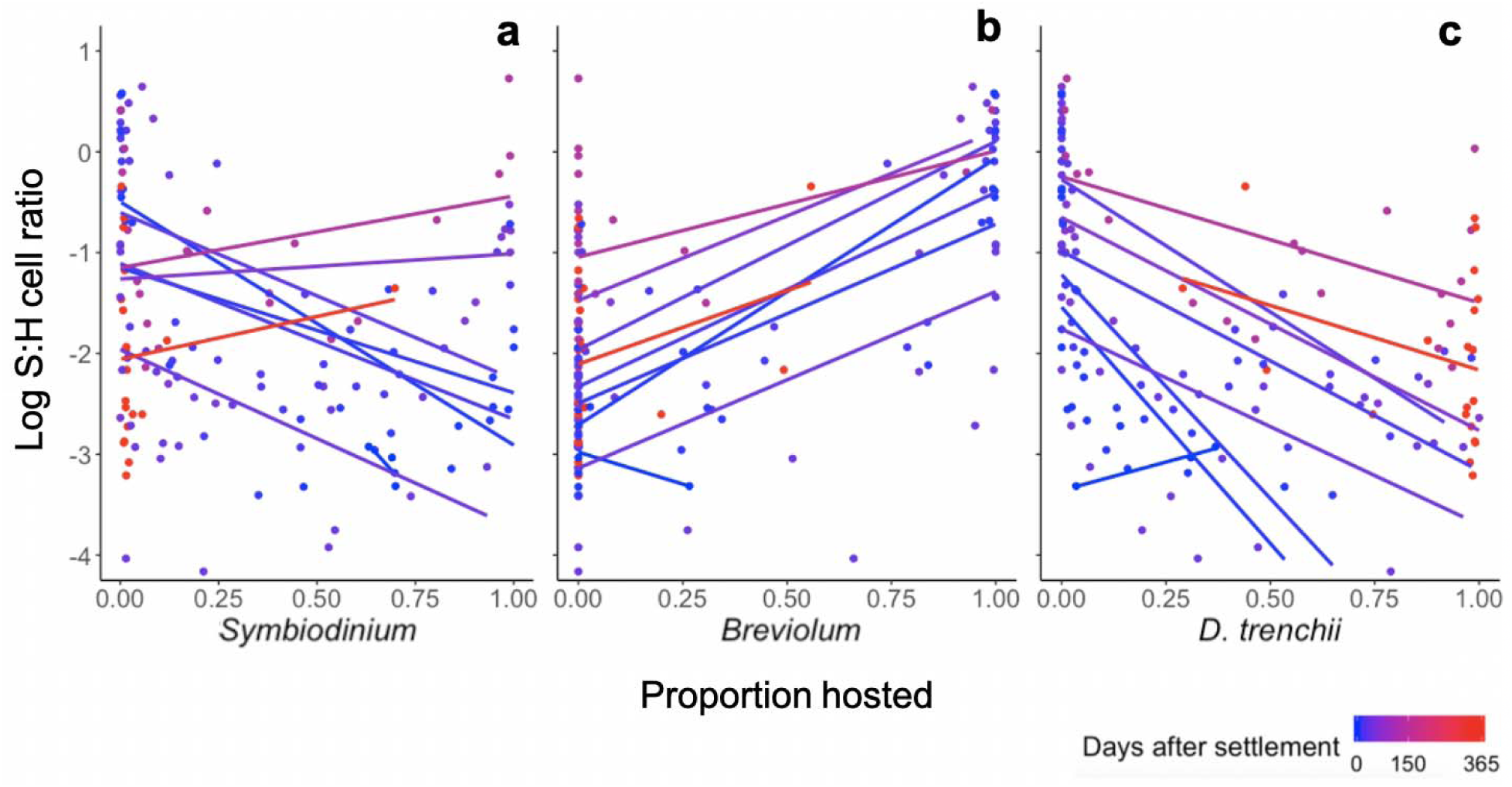
Symbiodiniaceae density varies with community composition in recruits. Log S:H cell ratio shows (a) no relationship with proportion of *Symbiodinium*, (b) positive correlation with proportion of *Breviolum* (p < 0.001), and (c) negative correlation with proportion of *D. trenchii* (p < 0.001). Points were jittered to minimize overlap.

### Polyp area

After 5 months, polyp size varied significantly with Symbiodiniaceae community composition. Mean polyp area increased with proportion *Breviolum* (p = 0.005, Fig. 6b) and decreased with increasing proportion of *D. trenchii* (p = 0.006) (Fig. 6c). There was no significant relationship between polyp size and previous temperature treatment or proportion of *Symbiodinium* (Fig. 6a).

**Figure 6:**
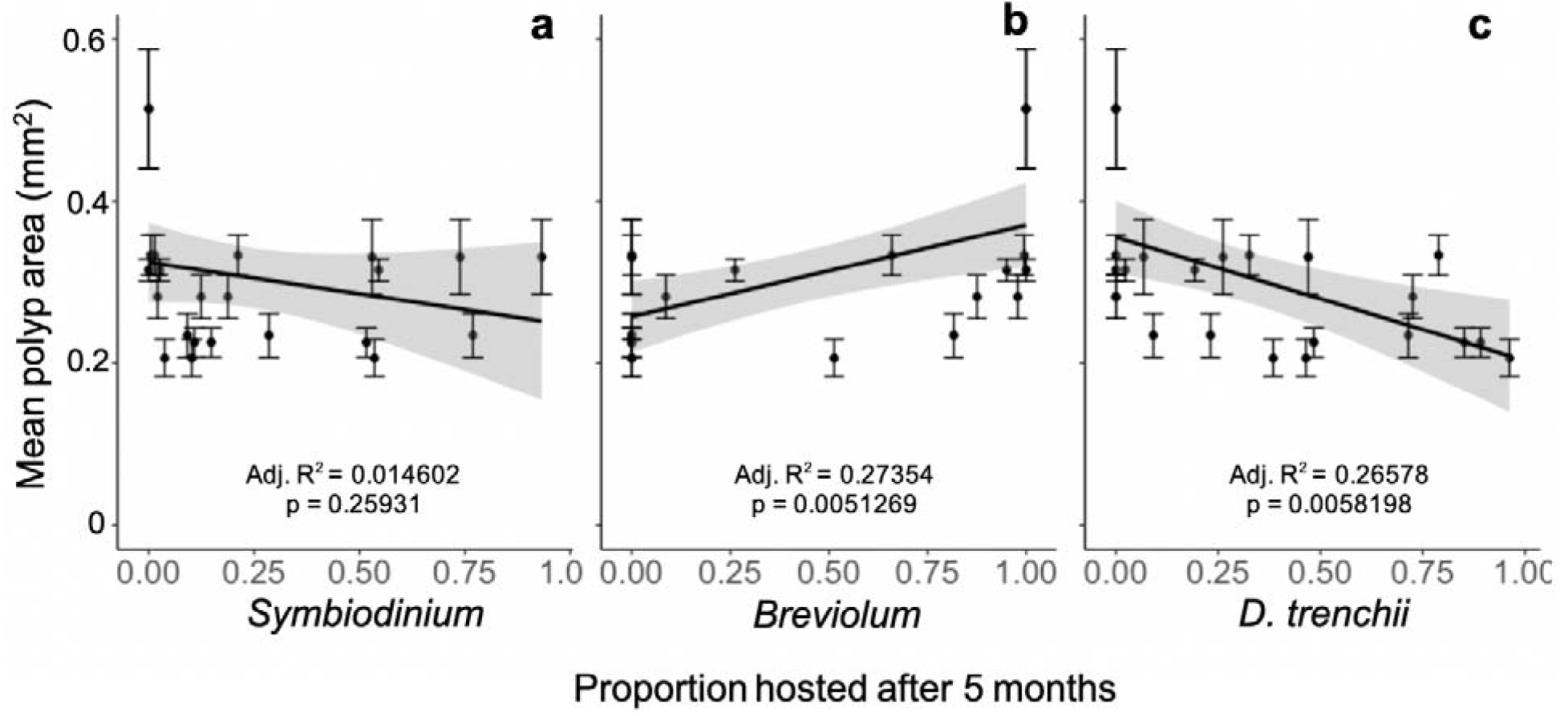
Correlation of polyp area with proportion of (a) *Symbiodinium*, (b) *Breviolum*, and (c) *D. trenchii*. Each point represents the mean size of polyps on one plug. Error bars represent +/- SEM. Points were jittered to prevent overlap.

### Experiment 2: Heat stress trial

During the heat stress trial, both previous heat exposure and symbiont community impacted recruit survivorship. Recruits previously reared at 31°C survived over twice as long at 34°C as those reared at 29°C, independent of their symbionts (p < 0.001) (Fig. 7a). In addition, recruits hosting high proportions of *D. trenchii* at the start of the trial survived over twice as long at 34°C as recruits with low proportions of *D. trenchii* and 50% longer than recruits with intermediate proportions of *D. trenchii* (p < 0.001) (Fig. 7b). Conversely, recruits with high proportions of *Symbiodinium* and *Breviolum* experienced reduced survivorship (Fig. S1). Survival probability during heat exposure was positively correlated with proportion of *D. trenchii* (p = 0.006) and negatively correlated with proportion of *Symbiodinium* (p = 0.022), but not correlated with *Breviolum* (p = 0.174).

**Figure 7:**
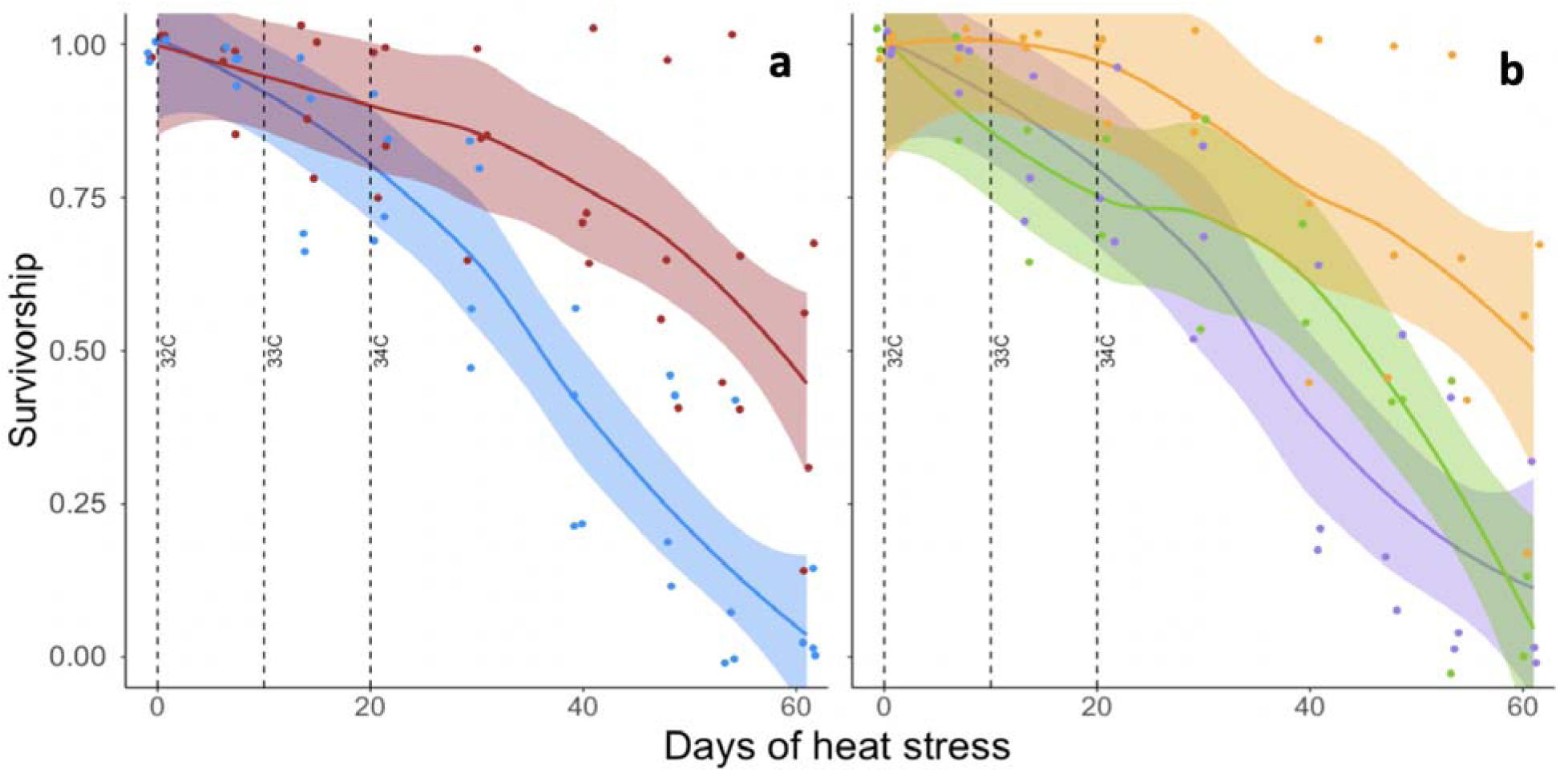
Heat stress trial revealed highest survivorship in recruits a) previously reared at 31°C and b) hosting a high proportion of *D. trenchii*. In panel (a), red points indicate recruits previously reared at 31°C and blue points indicate those reared at 29°C. In panel (b), orange points indicate recruits hosting high proportions of *D. trenchii* at the start of heat stress (>0.75), green indicates intermediate proportions (0.25 – 0.75), and purple indicates low proportions (<0.25). Points were jittered to minimize overlap. Lines indicate smoothed data (method = “loess”), and shaded areas indicate +/- SEM.

Similarly, both previous heat exposure and proportion of *D. trenchii* influenced bleaching resistance. During heat stress, recruits previously reared at 31°C and/or hosting high proportions of *D. trenchii* at the start of the trial maintained symbiosis longer than those reared at 29°C and/or hosting intermediate or low proportions of *D. trenchii* (Fig. S2, S3).

## Discussion

### Manipulating symbiosis establishment in recruits

This study tested a novel, scalable approach for introducing thermotolerant *Durusdinium trenchii* into coral hosts at a key stage in their life history. Our results indicate that temperature and/or symbionts in neighboring colonies can affect: (1) the rate of infection in *O. faveolata* recruits, (2) the identity of Symbiodiniaceae acquired during initial symbiosis establishment, and (3) the subsequent thermal tolerance of the recruits.

In experiment 1, temperature impacted the onset of symbiosis while nearby donor colonies influenced initial Symbiodiniaceae community composition. Infection was delayed in recruits reared at 31°C (a treatment representing only mild temperature stress), suggesting that even minimal heat exposure can slow algal acquisition in recruits by ∼5-20 days (Fig. 3). Abrego et al. (2012) similarly found that *Acropora tenuis* and *A. millepora* recruits at 31°C acquired symbionts much later than those at 28°C. Delayed infection could prove detrimental, particularly in the field, because juvenile corals rely on the fixed carbon their endosymbionts provide to grow and surpass size-escape thresholds (Doropoulos et al. 2012; Suzuki et al. 2013; Chamberland et al. 2017). Although we hypothesized that elevated temperature would favor acquisition of *D. trenchii* (Cunning et al. 2015a, Abrego et al. 2012), we found no significant effect of temperature on the algal species hosted, suggesting that availability outweighs temperature in determining the dominant symbiont type of *O. faveolata* recruits. However, it is also possible that 31°C was not hot enough to favor *D. trenchii* acquisition in recruits from Key Largo, FL.

In all treatments, recruits acquired *Symbiodinium, Breviolum*, and *D. trenchii*. This result is consistent with studies of other coral species, which have found that recruits can initially host a variety of different algal partners (Coffroth et al. 2001; Little et al. 2004; Mieog et al. 2009; Cumbo et al. 2018). Field surveys have found that *O. faveolata* adults associate with *Symbiodinium, Breviolum, Cladocopium*, and *Durusdinium* throughout their range and even within the same colony (Rowan & Knowlton 1995, Rowan et al. 1997; Kemp et al. 2015). In this study, the presence of *D. trenchii*-dominated *Siderastrea siderea* colonies enhanced the proportion of *D. trenchii* two-to three-fold in *O. faveolata* recruits. Nearby corals may influence symbiont availability by discharging algal cells, which persist in the sediments and water column and increase the rate of acquisition by newly-settled recruits (Coffroth et al. 2006; Nitschke et al. 2016; Ali et al. 2019). *D. trenchii* has been shown to be highly infectious in aposymbiotic recruits compared with other symbionts, although its abundance generally declines during ontogeny (Abrego et al. 2009; Pollock et al. 2017).

No recruits acquired detectable levels of *Cladocopium*, including those in aquaria with *Cladocopium* donor fragments. In *O. faveolata, Cladocopium* are generally found under relatively low irradiance around a colony’s base or on reefs deeper than 3 m (Rowan & Knowlton 1995; Kemp et al. 2015). Irradiance in our experiments (150–300 micromoles quanta m−2 s−1) may have resembled a shallow reef, where *O. faveolata* hosts primarily *Symbiodinium* and *Breviolum* (particularly B1) (Rowan et al. 1997; LaJeunesse 2002; Thornhill et al. 2009). Alternatively, since aposymbiotic cnidarians acquire “homologous” (native) symbionts more readily than other types (Weis et al. 2001, Mellas et al. 2014), the *Cladocopium* strain(s) available may have been “heterologous” to *O. faveolata*. Throughout its range, *M. cavernosa* typically hosts C1 and C3, whereas *O. faveolata* hosts C3, C7, and/or C12 (Thornhill et al. 2009; Kemp et al. 2015; Hauff et al. 2016). Fewer recruits in *Cladocopium* aquaria were infected after 43 days compared with those in *D. trenchii* aquaria (Fig. 3b), indicating a possible shortage of compatible Symbiodiniaceae.

Since *Symbiodinium* and *Breviolum* were detected in recruits but not in donor colonies, these algae must have been present at low abundance, either in the donor tissue, unsampled parts of donor fragments, the FSW, or the biofilm on the settlement plugs. As such, *Symbiodinium* and *Breviolum* were likely present at much lower densities than *Cladocopium* and *D. trenchii*, which were presumably shed by donor fragments during the experiment to regulate algal density in their tissues (Fitt et al. 2001). Therefore, the differential acquisition of *Symbiodinium* and *Breviolum* despite their relative scarcity suggests a degree of symbiont specificity in *O. faveolata* recruits under these conditions.

### Heat tolerance in *O. faveolata* recruits hosting *D. trenchii*

Recruits initially reared at 31°C survived significantly longer during the subsequent heat stress trial than those reared at 29°C, regardless of dominant symbiont type (Fig 7a), suggesting that pre-exposure to mild thermal stress may increase a juvenile’s ability to withstand future, more severe heat waves. It should be noted that in nature, coral recruits are unlikely to experience heat stress at 5 months post-settlement. Instead, since most broadcast spawning corals are born during the warmest time of year, they might more plausibly face heat stress either 0 - 3 months or one year after settlement. However, our results show that recruits can be preconditioned to heat, and that this effect may last at least 3 months after heat exposure ends. Future studies should test how long recruits retain thermal tolerance after pre-exposure.

Symbiodiniaceae community composition also impacted survivorship under heat stress. Recruits that started the trial with the highest proportion of *D. trenchii* (> 0.75) survived over twice as long at 34°C as recruits with low proportions of *D. trenchii* (< 0.25) and 50% longer than recruits with intermediate proportions (0.25 – 0.75) for a similar decline in survivorship (Fig. 7b). This agrees with previous studies reporting increased heat tolerance and bleaching resistance in adult *O. faveolata* hosting *D. trenchii* (Kemp et al. 2014; Cunning et al. 2018; Manzello et al. 2018). Acroporid juveniles from the Great Barrier Reef also survived better under thermal stress when hosting *D. trenchii* compared with other symbionts (Mieog et al. 2009; Abrego et al. 2012; Quigley et al. 2020; but see Abrego et al. 2008). Together, these findings suggest that *D. trenchii* may enhance coral survival during vulnerable early life stages in the face marine heatwaves that can often coincide with, or shortly follow, coral spawning events.

### Trade-offs of hosting *D. trenchii*

Symbiodiniaceae cell density varied with algal community composition in *O. faveolata* recruits. At all time points, log S:H cell ratio was negatively correlated with the proportion of *D. trenchii* (Fig. 5c) and positively correlated with the proportion of *Breviolum* (Fig. 5b). Fewer symbionts per host cell could indicate slower colonization or proliferation rates for *D. trenchii*, differences in cell size between symbiont species (Jones & Yellowlees 1997; LaJeunesse et al. 2018), or a host-regulated reduction in symbiont density when hosting *D. trenchii*. This low cell S:H cell ratio may also help explain the heat tolerance observed in recruits hosting *D. trenchii*, since high Symbiodiniaceae density can increase bleaching susceptibility (Cunning & Baker 2013).

We observed a significant negative correlation between the proportion of *D. trenchii* and polyp size after 5 months, consistent with previous studies reporting reduced growth in juvenile and adult corals hosting *D. trenchii* (Little et al. 2004; Cantin et al, 2009; Jones & Berkelmans 2010; Pettay et al. 2015; but see Yuyama & Higuchi 2014; Quigley et al. 2020). For instance, Little et al. (2004) found that acroporid recruits infected with *Cladocopium* grew 2-3 times faster than those hosting *Durusdinium*. Under ambient, non-stressful conditions, *D. trenchii* is less photochemically efficient than *Breviolum* and *Cladocopium* (Cunning et al. 2018) and thus may have less photosynthate available for the coral host. Although dominant Symbiodiniaceae type explained up to 30% of variation in polyp size after 5 months, algal communities detected at the time of photographing and sampling provided only a “snapshot” assessment and do not necessarily represent previous communities that may have contributed to growth earlier in ontogeny. However, due to their small size (<1 mm in diameter), it was impossible to sample a recruit’s symbiont community without sacrificing it. Consequently, some of the unexplained variation in polyp size could be due to changes in symbiont communities over time. Algal density could also impact growth, because fewer Symbiodiniaceae cells in coral tissues may translate to less carbon produced and delivered to the host. Thus, the smaller size of recruits hosting *D. trenchii* may result from their low symbiont density, and not necessarily from these symbionts being individually stingy or “selfish” (Stat & Gates 2011).

Growth rates can help determine winners and losers in coral reef ecosystems, which are highly competitive for space and light. Like many organisms, corals attain some refuge in size whereby predators and competitors are less likely to harm larger individuals. Field studies of *Pocillopora damicornis* and *Siderastrea radians* juveniles reported significant increases in survivorship with colony size (Raymundo & Maypa 2004; Vermeij & Sandin 2008). Combined with environmental stressors such as ocean acidification that already compromise coral growth (Hoegh-Guldberg et al. 2007; Doropoulos et al. 2012), hosting *D. trenchii* could further prolong the vulnerability that comes with small size and prevent population recovery (Ortiz et al., 2013). With these potential trade-offs, it will be important to take many factors into account when assessing the net risks versus benefits of *D. trenchii* in enhancing coral restoration strategies.

### Implications for reef restoration and resilience

In recent years, researchers, practitioners, and managers have recognized the importance of sexually-derived coral stock to expanding the genetic diversity and spatial scale of reef restoration efforts (Baums et al. 2019, Randall et al., 2020). However, strategies for maximizing the early post-settlement survival of reared and outplanted juveniles need to be advanced. Specifically, efforts to: (1) reduce the risk of predation or competition, (2) promote early, beneficial associations with Symbiodiniaceae, and (3) enhance stress tolerance prior to outplanting can minimize juvenile mortality and strengthen the impact of restoration efforts (Quigley et al. 2018).

Since *O. faveolata* (and most Caribbean reef-building corals) spawn during the warmest time of year, recruits are likely to experience heat stress during the first few months of their lives, a threat that will only intensify with continued climate change. As such, boosting the thermal resilience of new generations of corals should be a priority of reef restoration efforts. Since *D. trenchii* may enable recruits to resist heat stress, restoration practitioners concerned about bleaching risk may benefit by rearing recruits near *D. trenchii* donors (*in situ* or *ex situ*) to increase uptake of this algae. However, experiments to manipulate symbioses may only be feasible in certain species. While some coral juveniles can acquire diverse Symbiodiniaceae during early ontogeny (Little et al. 2004; Abrego et al. 2009; Cumbo et al. 2013; McIlroy & Coffroth 2017), others exhibit genetically-determined symbiont preferences (Weis et al. 2001; Poland & Coffroth 2017; Quigley et al. 2017b). Similarly, the relative benefit of hosting *D. trenchii* likely varies among species (Cunning et al. 2018). Therefore, restoration practitioners should test how *D. trenchii* impacts the particular coral(s) they work with before incorporating it into their efforts.

Given its associated physiological trade-offs, there is a risk that *D. trenchii* may reduce corals’ competitive ability and prolong recovery following disturbances (Ortiz et al., 2013). Nevertheless, since a majority of the world’s reefs are projected to experience annual severe bleaching by mid-century (van Hooidonk et al. 2014), coral persistence may soon depend more on heat tolerance than growth. Under elevated temperature, *D. trenchii* confers a photochemical advantage to its hosts and relative growth trade-offs may decrease or even disappear at progressively higher temperatures (Cunning et al. 2015b; Cunning et al. 2018). Thus, under a ‘new normal’ of repeated or chronic heat stress, *D. trenchii* may increase coral survival without compromising growth relative to other symbiont genera. Stakeholders should therefore consider climate projections when deciding if *D. trenchii* may help or harm the reefs they manage in the long-term.

Going forward, it will be important to determine whether recruits can be primed with *D. trenchii in-situ* as part of existing restoration pipelines. Without the constraints of laboratory space, field methods to boost *D. trenchii* uptake could help practitioners rear large numbers of thermally-tolerant coral juveniles for outplanting. However, even if *D. trenchii* proves advantageous during thermal anomalies, it may be lost from corals over time in the absence of heat stress (Thornhill et al. 2006; LaJeunesse et al. 2009). Therefore, future studies should examine the longevity of manipulated symbiont communities in outplanted recruits, and identify conditions that promote *D. trenchii* dominance and maintain its benefits for hosts.

Finally, our finding that nearby colonies enhance *D. trenchii* uptake in recruits may inform the potential for symbiont community feedbacks within and between generations. While it was historically present at low abundances in some corals and locations, *D. trenchii* has rarely served as the dominant, preferred symbiont (e.g., LaJeunesse et al. 2002) unless colonies experienced environmental stress or extremes (Baker et al. 2004; Fabricius et al. 2004; Kennedy et al. 2015; Silverstein et al. 2015). However, warming oceans and recurring bleaching events may favor *D. trenchii* at the ecosystem level because it colonizes newly-settled recruits and adults recovering from bleaching (Nitschke et al. 2016; Boulotte et al. 2016), and colonies that cannot shift to environmentally-appropriate partnerships perish (LaJeunesse et al. 2010; Grottoli et al. 2014).

It remains uncertain whether the proliferation of *D. trenchii* on coral reefs will promote resilience or hinder recovery. As studies continue to disentangle the relationships between Symbiodiniaceae identity, coral physiology, and environmental variability, we can begin to predict context-dependent trade-offs within the coral holobiont and use our findings to inform reef restoration. If early infection with *D. trenchii* increases thermal tolerance in coral juveniles without severely compromising other aspects of fitness, practitioners may choose to rear them near *D. trenchii*-dominated adult colonies, boosting their resilience before outplanting them onto the reef.

## Acknowledgements

Fieldwork was conducted under permit FKNMS-2016-047-A1. The authors thank A. Peterson and A. Bright for support during coral spawning dives, and for rearing *O. faveolata* larvae prior to settlement. OW thanks M. Schmale for use of his dissecting microscope to sample recruits, R. Cunning for support with data analysis, and R. van Hooidonk for calculating sea surface temperature MMM’s using Pathfinder 5.0 and Coral Reef Watch. Funding sources: CIMAS Fellowship and B. Kirtman, NOAA Task III (to M. Johnson and A. Baker)

## Conflict of Interest Statement

On behalf of all authors, the corresponding author states that there is no conflict of interest.

## Notes

### Competing Interest Statement

The authors have declared no competing interest.

