## Supplementary Information for "Neighboring colonies influence uptake of thermotolerant endosymbionts in threatened Caribbean coral recruits"

**Neighboring colonies influence algal symbiont community structure in threatened Caribbean coral recruits**

**Supplementary information**

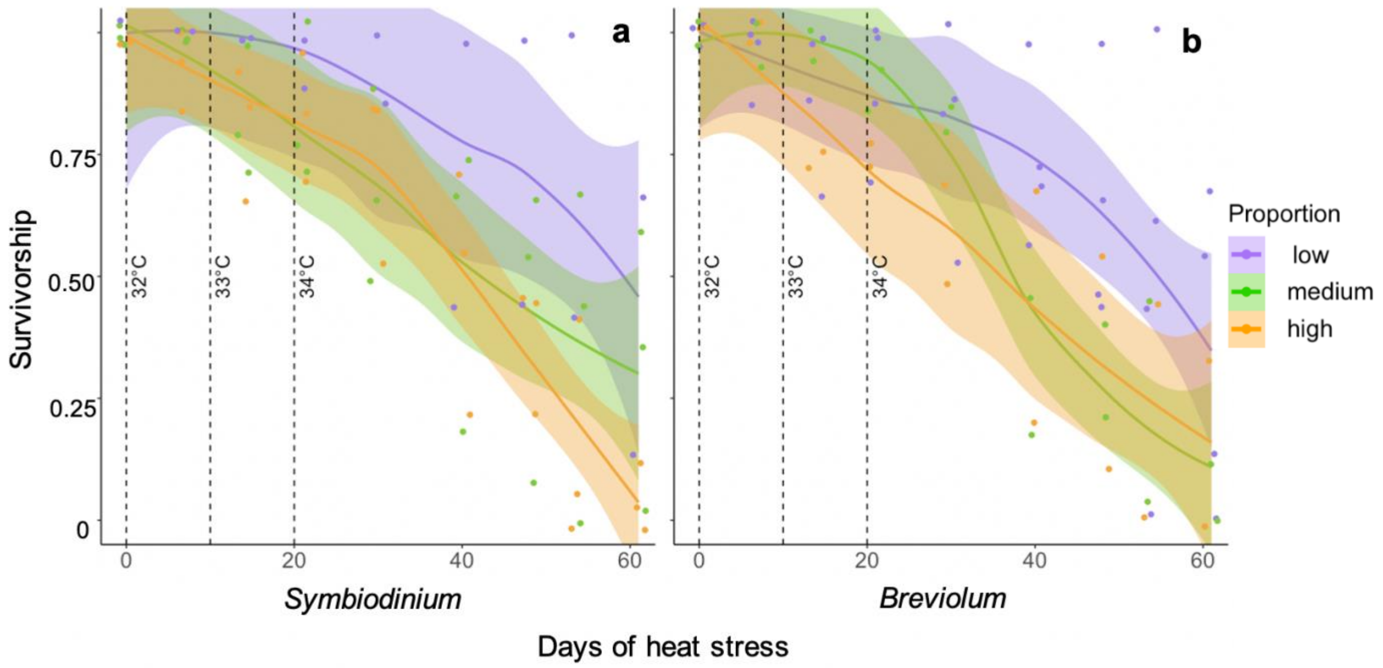

**Figure S1: Recruits with high proportions of a) *Symbiodinium* and b) *Breviolum* survived half as long at 34°C compared the recruits with low proportions of these symbionts.** In panel a), “high” proportion of *Symbiodinium* refers to 0.4 – 0.7, “intermediate” to 0.1 – 0.4, and “low” to 0 – 0.1. In panel b), “high” proportion of *Breviolum* refers to 0.1 – 0.4, “intermediate” to 0 – 0.1, and “low” to 0. Points were jittered around each time point to improve visualization of data. Lines indicate smoothed data (method = “loess”), and shaded areas indicate +/- SEM

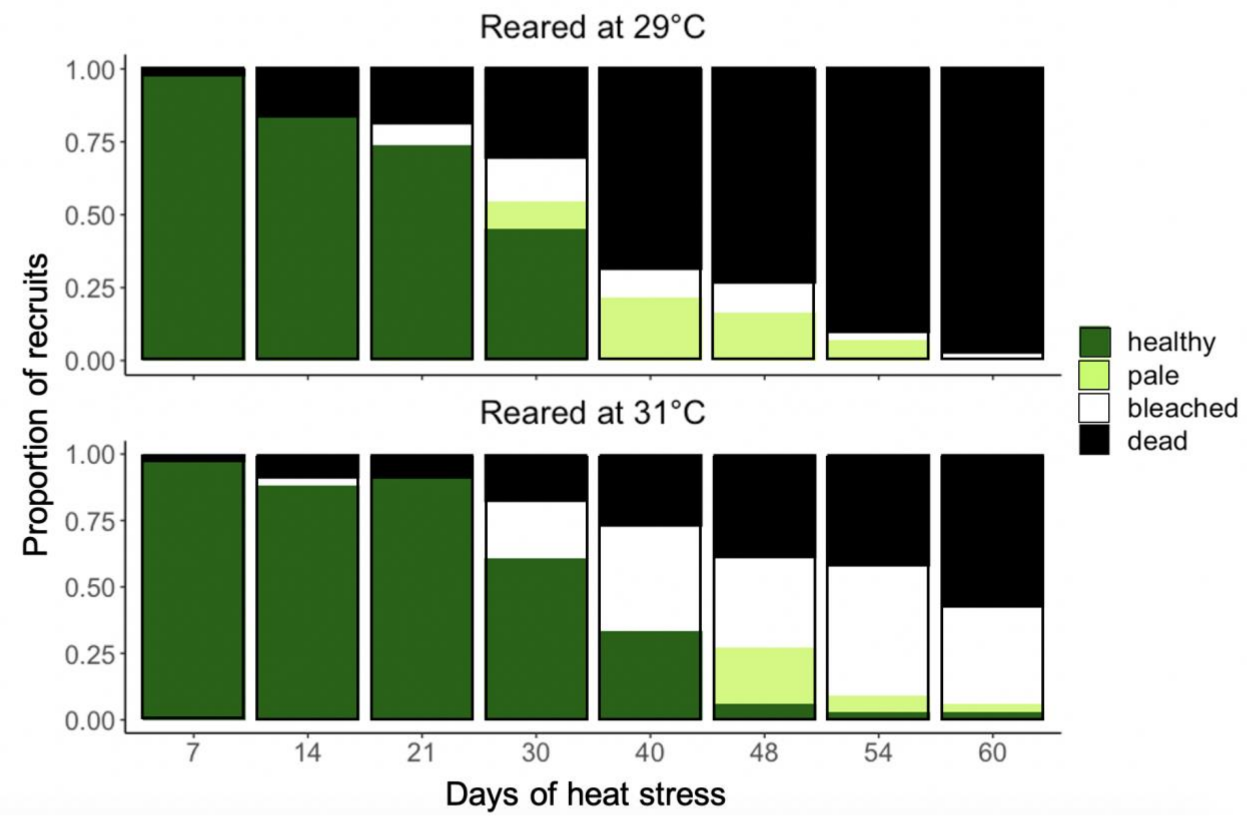


**Figure S2: Health status during heat stress of 5-month-old *O. faveolata* recruits a) previously reared at 29°C and b) previously reared at 31°C**. The proportion of recruits maintaining symbionts (“healthy”) was significantly higher among recruits previously reared at 31°C compared to recruits previously reared at 29°C (ANOVA, GLM, p = 0.02). There was a significant interaction between days of heat stress, rearing temperature, and proportion of “healthy” recruits (p = 0.001)

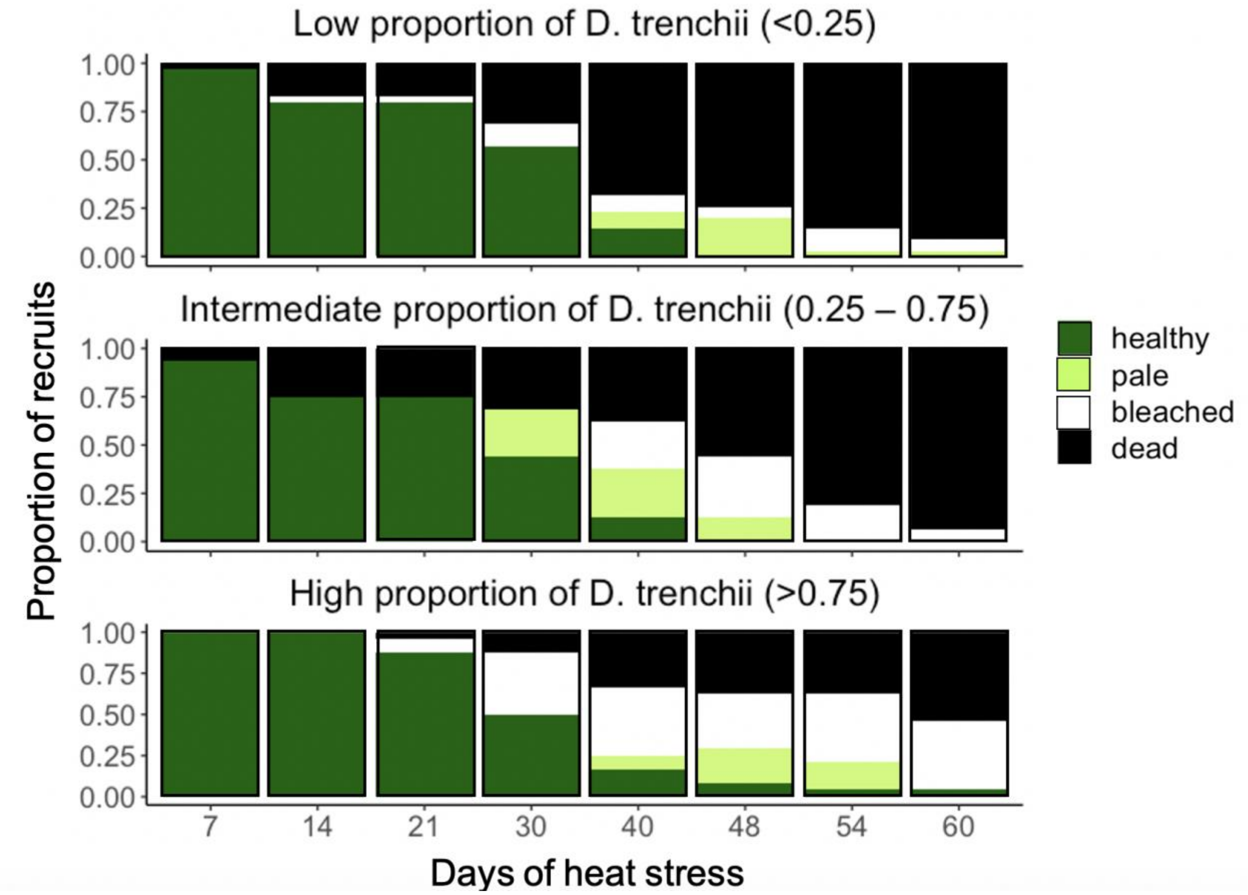


**Figure S3: Health status during heat stress of 5-month-old *O. faveolata* recruits hosting a) a low proportion of *D. trenchii* (<0.25), b) an intermediate proportion of *D. trenchii* (0.25 – 0.75), and c) a high proportion of *D. trenchii* (>0.75)**. According to ANOVA and generalized linear models, there was a significant interaction between days of heat stress, proportion of *D. trenchii* hosted by recruits, and proportion of “healthy” recruits ($p = 0.002$).
